# CDC20 determines the sensitivity to spindle assembly checkpoint (SAC) inhibitors

**DOI:** 10.1101/2023.12.21.572559

**Authors:** Siqi Zheng, Linoy Raz, Lin Zhou, Yael Cohen-Sharir, Ruifang Tian, Rene Wardenaar, Mathilde Broekhuis, Maria Suarez Peredo Rodriguez, Soraya Wobben, Anouk van den Brink, Petra Bakker, Floris Foijer, Uri-Ben David

## Abstract

Spindle assembly checkpoint (SAC) inhibitors are a recently developed class of drugs that perturb the regulation of chromosome segregation during division, induce chromosomal instability (CIN), and eventually lead to cell death. While they are currently in clinical trials for solid cancers, biomarkers to predict the response to SAC inhibitors are still lacking. We recently reported that aneuploid cancer cells are preferentially sensitive to SAC inhibition. Here, we investigated the molecular determinants of the response to SAC inhibition that underlies the differential sensitivity of aneuploid cells to these drugs. We found that this response was largely driven by the expression of CDC20, a main activator of the anaphase-promoting complex (APC/C), rather than by APC/C itself. Mechanistically, we discovered that CDC20 depletion prolonged metaphase duration, diminished mitotic errors, and reduced sensitivity to SAC inhibition. Aneuploid cells expressed high levels of CDC20 and experienced shorter metaphases and multiple mitotic errors, resulting in increased long-term sensitivity to SAC inhibition. Our findings propose high CDC20 expression as a favorable biomarker for SAC inhibition therapy and as an aneuploidy-induced therapeutic vulnerability.

## Introduction

The spindle assembly checkpoint (SAC), also known as the mitotic checkpoint, is a key cell cycle regulator that ensures the fidelity of chromosome segregation during mitosis. In the presence of abnormal mitotic spindles that will lead to errors in chromosome segregation, the SAC is recruited and arrests cell division, allowing time for error correction and restoration of normal division. The SAC is composed of a series of proteins that recruit an effector complex, the mitotic checkpoint complex (MCC). The MCC, which is composed of BUB1B, MAD2L1 and BUB3, prevents progression into anaphase by sequestering CDC20, the main activator of the anaphase-promoting complex (APC/C) (Curtis et al, 2020; Kops et al, 2005). When unbound by the MCC, CDC20 forms a complex with the E3 ubiquitin ligase APC/C, and the complex tags various substrates for degradation by ubiquitination. Notable CDC20-APC/C substrates are Securin, whose degradation allows sister chromatid separation; Cyclin B1, whose degradation also promotes mitosis culmination; and CDC20 itself, which is negatively regulated by the complex (Musacchio & Salmon, 2007). However, when the SAC is active, the MCC binds CDC20 and progression into anaphase is blocked, allowing time for error correction and restoration of normal division (Musacchio & Salmon, 2007). Nonetheless, prolonged cell cycle arrest or failure to divide normally will eventually lead either to cell death or to mitotic slippage (Sinha et al, 2019, Rossio et al, 2010) – cell division despite the presence of an altered spindle and an active SAC – which would lead to chromosome missegregation and aneuploidization.

SAC inhibitors are drugs that disable the SAC, enabling cells to complete erroneous divisions and causing the equivalent of mitotic slippage. Consequently, these drugs induce chromosomal instability and lead to the acquisition of aberrant and unfit karyotypes in the treated cells (He et al, 2018, Mason et al, 2017, Kawakami et al, 2019). We recently showed that SAC inhibition is more detrimental for aneuploid cells than for diploid cells (Cohen-Sharir et al, 2021). SAC inhibitor drugs – and specifically, MPS1 inhibitors – are currently in multiple clinical trials for the treatment of solid cancers (NCT02792465, NCT03568422, NCT05251714), either alone or in combination with microtubule-disrupting agents such as paclitaxel. Despite the emerging clinical utility of these drugs, no biomarker predicting patient response has yet been confirmed, and the molecular mechanism behind the response to these drugs is only partially understood. Several papers suggested that low expression levels or dysfunction of the APC/C are associated with resistance to SAC inhibition (Sansregret et al, 2017, Thu et al, 2018, Wild et al, 2016). However, in this work we refine the proposed mechanism underlying the response to SAC inhibition, suggesting that CDC20 regulates this response. We further propose that CDC20 underlies the differential sensitivity of aneuploid cells to these drugs.

## Results

### Cdc20 is strongly associated with the response to SAC inhibition

To identify genes and pathways that are involved in the response to SAC inhibition, we performed two independent CRISPR-Cas9 screens in NIH-3T3 mouse fibroblasts, looking for genes whose depletion promoted cell proliferation under SAC inhibition using the Mps1 inhibitor reversine, as shown in **Figure 1A** and Supplementary Fig. 1A,B. Comprehensive lists of the resultant candidate genes can be found in Supplementary Table 1. Searching for cellular pathways enriched in the top ranking 10% of candidate genes from both CRISPR screens, we found that three out of ten pathways were directly related to anaphase promotion by the APC/C, and a fourth pathway contained the APC/C complex genes among other nuclear ubiquitin-ligases (**Figure 1B**). Interestingly, the top significant gene whose loss conferred resistance to SAC inhibition in both screens was Cdc20, the protein co-activator of the APC/C during mitosis (**Figure 1C**). While the core members of the APC/C were among the candidate genes, their effect was much weaker than that of Cdc20 in both of the screens (Supplementary Fig. 1C). To validate the role of Cdc20 levels in promoting resistance to SAC inhibition, we knocked down or knocked out Cdc20 in NIH-3T3 cells using shRNA or CRISPR, respectively (Supplementary Fig. 1D-F), and assessed sensitivity to the SAC inhibitor reversine (Santaguida *et al*, 2010). Indeed, Cdc20 depletion reduced sensitivity to SAC inhibition, evidenced by increased cell viability following exposure to the drug (**Figure 1D** and Supplementary Fig. 1G). Together, these findings suggest a strong association between Cdc20 expression and sensitivity to SAC inhibition, with Cdc20 depletion leading to increased resistance to these drugs.

**Figure 1.**
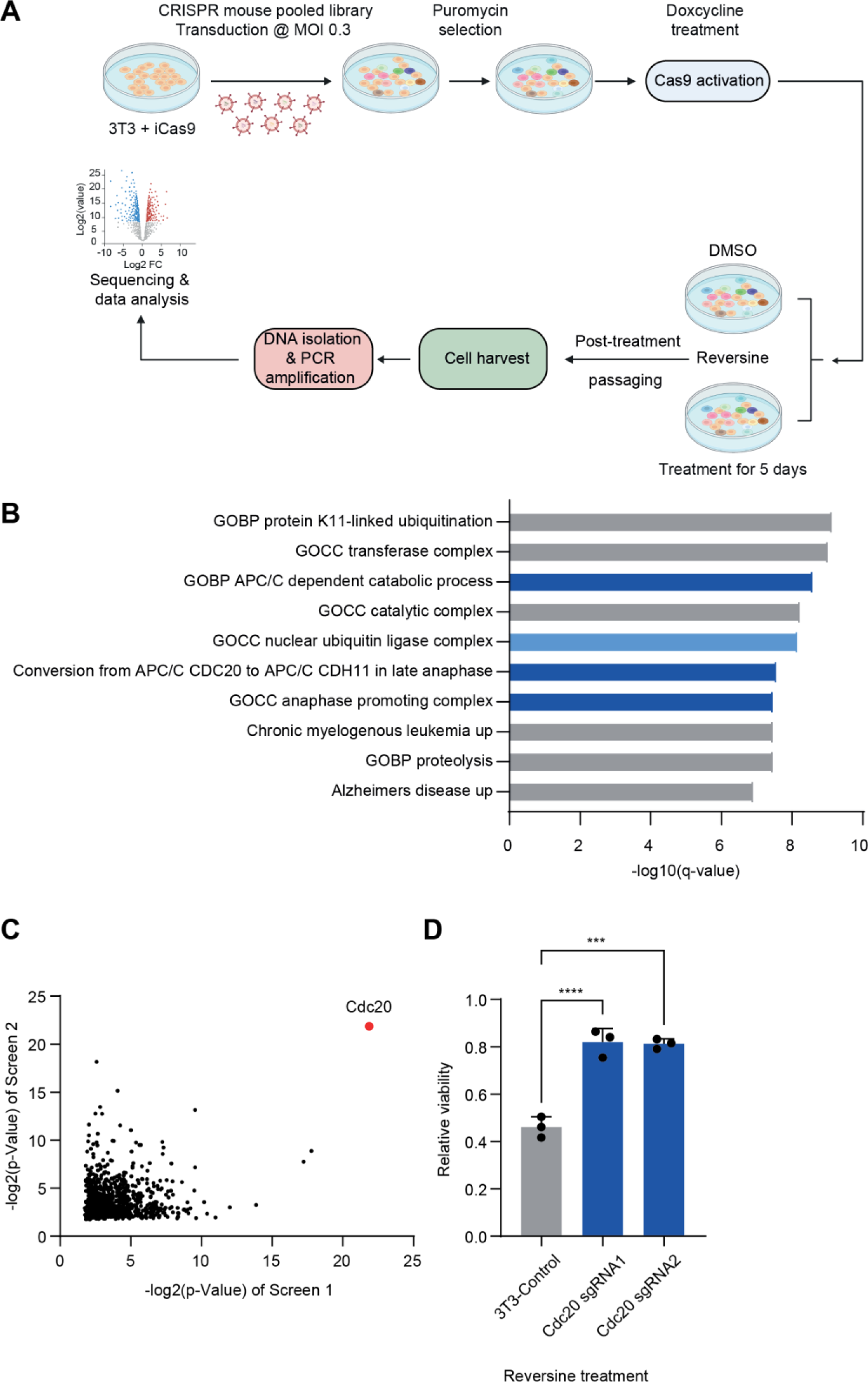
Cdc20 is strongly associated with the response to SAC inhibition. (A) Schematic overview of the CRISPR screens performed to identify genes and pathways involved in resistance to SAC inhibition. **(B)** Cellular pathways enriched in the top ranking 10% of genes in both CRISPR screens. **(C)** Correlation between the top ranked 25% of genes in both CRISPR screens, based on their statistical significance. Cdc20 (red) is the top hit in both screens. **(D)** Sensitivity of NIH-3T3 cells to 250nM of the SAC inhibitor reversine following Cdc20 knockout by CRISPR-Cas9. One-way ANOVA (N = 3, **, p-value <0.01, ***, p-value <0.001; ****, p-value < 0.0001)

### CDC20 expression, but not APC/C expression, predicts sensitivity to genetic and chemical SAC inhibition

Several studies suggested that resistance to SAC inhibition is associated with lower expression levels of the anaphase-promoting complex (APC/C) (Sansregret et al, 2017, Thu et al, 2018, Wild et al, 2016). However, our CRISPR screen suggested that it was Cdc20, rather than the APC/C core members, that determines the sensitivity to SAC inhibition. Indeed, while knockout and knockdown of Cdc20 reduced the sensitivity of NIH-3T3 cells to SAC inhibition (**Figure 1D** and **Supplementary Fig. 1G**), knockdown and knockout of a representative APC/C subunit, Anapc4, did not (**Supplementary Fig. 2A-E**).

To further explore the respective roles of CDC20 and the APC/C in the response to SAC inhibition, we analyzed genomic and transcriptomic data from over 1,700 human cancer cell lines from the cancer Dependency Map (Tsherniak et al, 2017). We used an APC/C transcriptional signature previously reported by Thu et al. to be associated with sensitivity to SAC inhibition (Thu et al, 2018) (**Supplementary Fig. 2F**) to assign an APC/C expression score to each cell line, and correlated the APC/C expression with sensitivity to genetic perturbation of the core SAC components MAD2L1 and BUB1B. Indeed, a lower APC/C expression score was correlated with lower sensitivity to genetic SAC perturbation (**Figure 2A,B**, left panels). However, we noticed that the gene signature used by Thu et al. to define the APC/C contained not only the core APC/C subunits but also CDC20, which acts as the main APC/C co-activator during mitosis. Intriguingly, this correlation was lost or severely weakened when we removed CDC20 and used a signature containing only the APC/C subunits (Yamano, 2019), (**Supplementary Fig. 2F** and **Figure 2A, B,** middle panels). In contrast, expression of CDC20 alone was strongly correlated with sensitivity to genetic SAC perturbation (**Figure 2A** **,B**, right panels). Next, we used the human cell line data to study the association between gene expression and sensitivity to chemical perturbation of the SAC. Here as well, increased CDC20 expression was correlated with increased sensitivity to SAC inhibitor drugs (Mps1 inhibitors MPI-0479605 and AZ3146), while APC/C subunit expression was not **(Figure 2C, D)**. Overall, the human cancer cell line data suggest that elevated levels of CDC20, but not of APC/C, expression is associated with increased sensitivity to both genetic and chemical disruption of the SAC.

**Figure 2.**
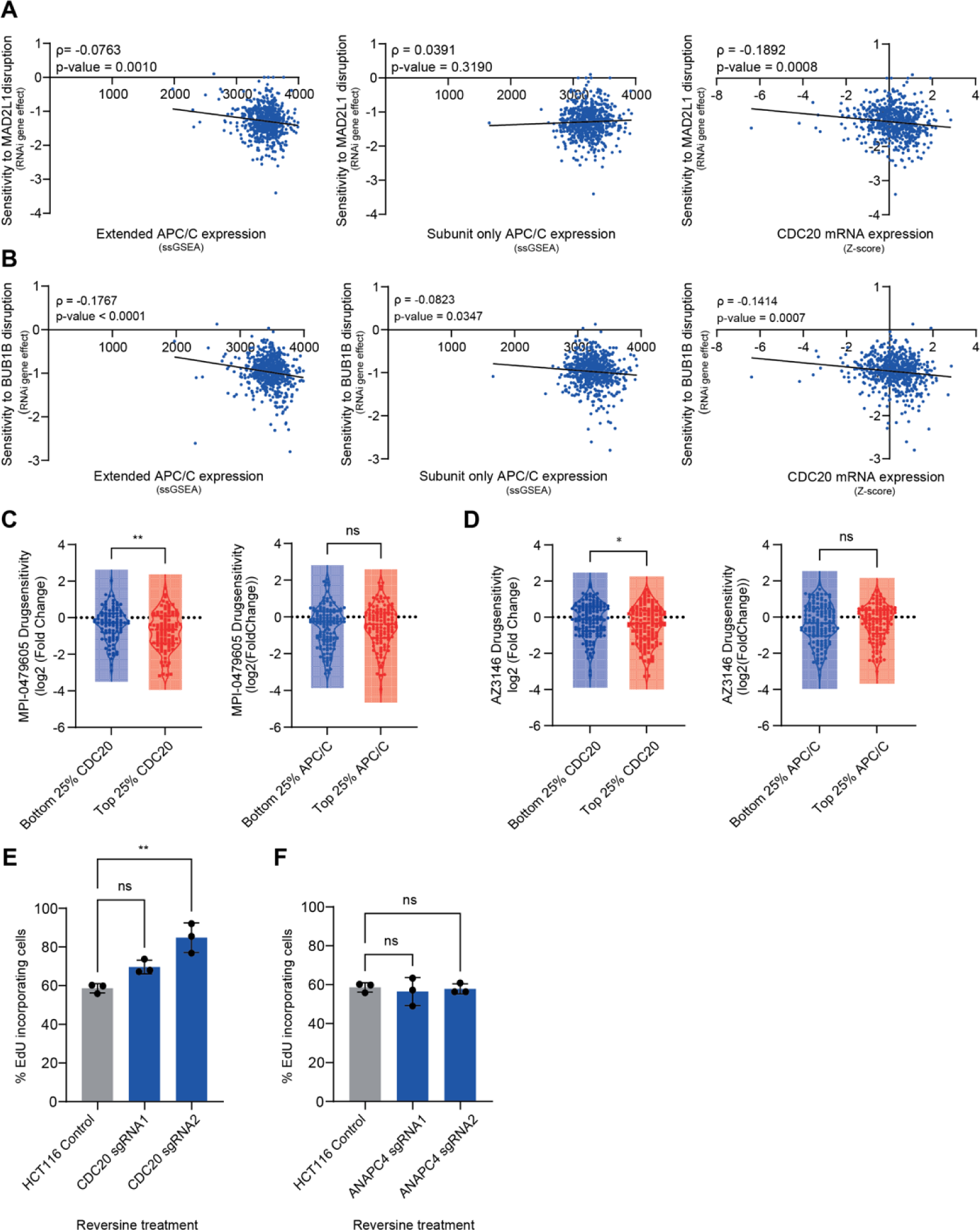
CDC20 expression, but not APC/C expression, predicts sensitivity to genetic and chemical SAC perturbation. **(A, B)** Correlation between mRNA expression of an extended APC/C signature (left), an APC/C signature without CDC20 (middle), or CDC20 alone (right) and sensitivity to genetic disruption of the core SAC components BUB1B **(A)** and MAD2L1 **(B)** in human cancer cell lines from the DepMap. The “APC/C subunit-only” signature contains the 14 core APC/C subunits (Yamano, 2019), while the “extended APC/C signature” described by Thu et al (Thu et al, 2018) contains three additional APC/C co-factors, including CDC20. The genes included in each signature are listed in **Supplementary Fig. 2A, B**. Shown are Spearman’s correlation rho and p values. (N=661 for MAD2L1 or BUB1B vs extended APC/C or subunit-only APC/C, N=662 for MAD2L1 or BUB1B vs extended APC/C or subunit-only APC/C; *, p-value<0.05; **, p-value<0.01) RNAi dependency scores were obtained from the Achilles genome-wide RNAi screen, DepMap 22Q2 (Tsherniak et al, 2017) **(C,D)** Comparison of the sensitivity to two chemical SAC inhibitors – MPI-0479605 **(C)** and AZ3146 **(D)** between cell lines in the top vs. bottom mRNA expression quartiles of CDC20 (left) and APC/C subunits (right). Drug sensitivity data were obtained from PRISM repurposing primary CRISPR screen, DepMap 19Q4 (Corsello et al, 2020). Two-sided T-test (N=260 for CDC20 vs MPI-0479605, N=198 for APC/C vs MPI-0479605, N=274 for CDC20 vs AZ3146, N=276 for APC/C vs AZ3146; *, p-value<0.05; **, p-value<0.01). **(E, F)** Percent of EdU-incorporating HCT116 cells following SAC inhibition (125nM reversine), with and without CDC20 **(E)** or ANAPC4 **(F)** knockout. CDC20 knockout increased the fraction of proliferating cells following drug treatment, while ANAPC4 knockout did not significantly change their incidence. One way ANOVA (N = 3; **, p-value < 0.01).

To validate these findings in the context of human cancer cells, we knocked out CDC20 and ANAPC4 in the human colon cancer cell line HCT116 (**Supplementary Fig. 2G, H**), and assessed cell survival and proliferation under SAC inhibition. Similar to the observations with the mouse cells, CDC20 depletion led to a substantial decrease in the cellular sensitivity of human cancer cells to SAC inhibition (**Figure 2E**). In contrast, ANAPC4 knockout did not exert a similar effect on drug response (**Figure 2F**). These results suggest that both in mouse and in human cells it is CDC20 expression, rather than expression of the APC/C core components, that mediate the cellular response to SAC inhibition.

### Increased CDC20 expression is associated with the preferential response of aneuploid cells to SAC inhibition

We recently found that aneuploid cells are more sensitive to SAC inhibition than diploid cells (Cohen-Sharir et al, 2021). We therefore wondered whether the expression levels of CDC20 differ between diploid and aneuploid cells, and if so, whether this underlies their differential drug sensitivity. We therefore compared CDC20 and APC/C mRNA expression levels between the top and bottom aneuploid quartiles of human cancer cell lines in the DepMap database (Tsherniak et al, 2017; Cohen-Sharir et al, 2021). We found that highly-aneuploid cell lines over-express CDC20, but under-express most of the core APC/C subunits (Yamano, 2019) in comparison to near-diploid cell lines (**Figure 3A**). This was surprising given that expression patterns of cell cycle genes are usually strongly correlated across cell lines (Fischer et al, 2022). We therefore examined highly-aneuploid and near-diploid cell lines separately and found that the mRNA and protein expression levels of CDC20 and the APC/C subunit ANAPC4 were indeed strongly correlated in the near-diploid cell lines, but this correlation was completely lost in highly-aneuploid cells (**Figure 3B**). Given the increased susceptibility of aneuploid cells to SAC inhibition in comparison to diploid cells (Cohen-Sharir et al, 2021), these results further support the notion that the cellular response to SAC inhibition is primarily determined by quantitative changes in CDC20 expression rather than by the expression of the APC/C subunits.

**Figure 3.**
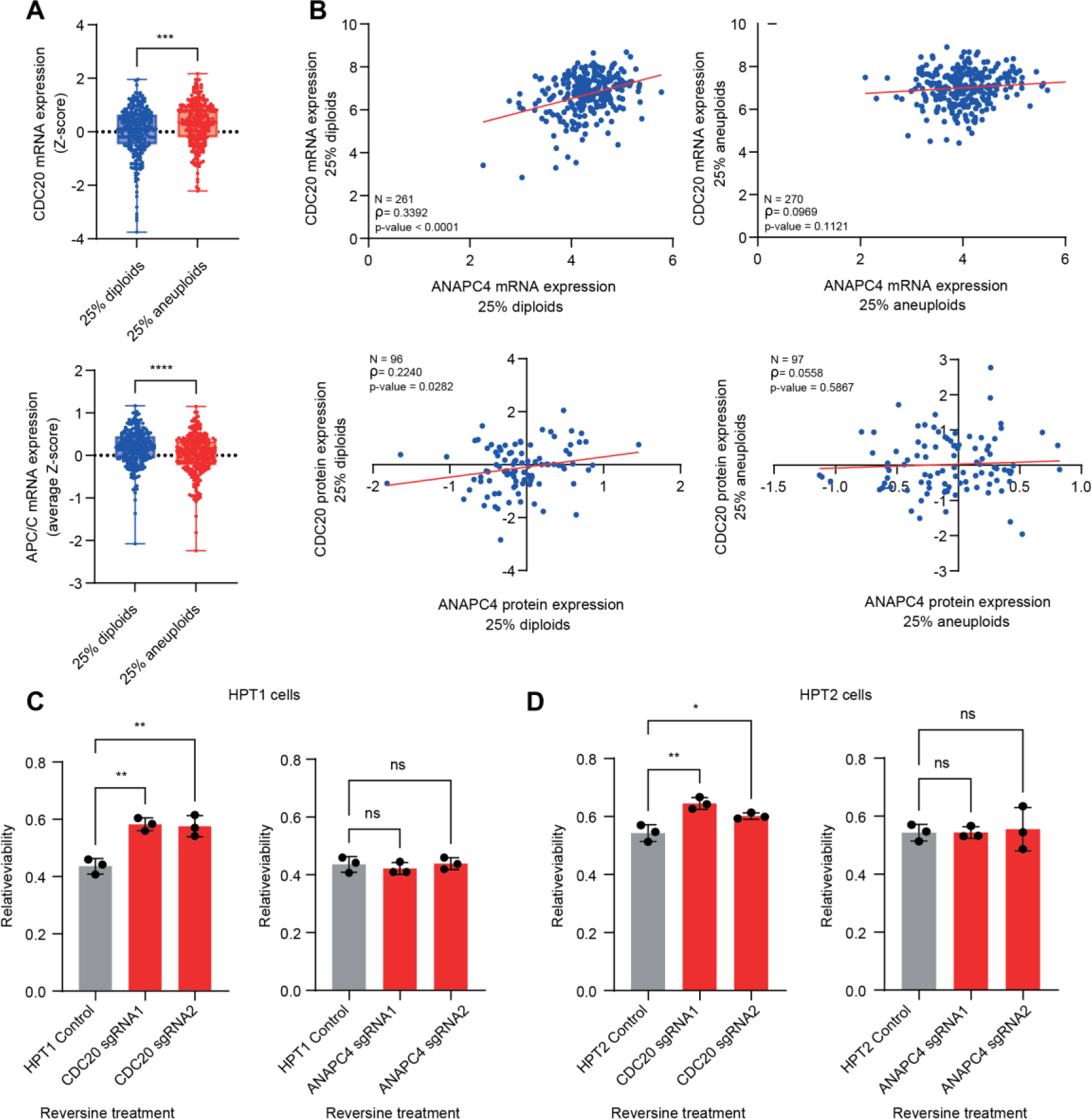
Increased CDC20 expression is associated with the preferential response of aneuploid cells to SAC inhibition. (**A-D**) Analysis of mRNA and protein expression levels in ∼1,000 human cancer cell lines. (**A**) Highly-aneuploid cancer cell lines express significantly higher mRNA levels of CDC20, but overall lower levels of the APC/C, compared to near-diploid cell lines. Two-sided T-test (N=531, ***, p-value<0.001; ****, p-value<0.0001). **(B)** Correlation between CDC20 and ANAPC4 expression in the top and bottom aneuploidy quartiles of cancer cell lines. Top panels: mRNA expression; bottom panels: protein expression. ANAPC4 was selected as a representative APC/C subunit with available proteomics data. CDC20 and ANAPC4 mRNA and protein expression levels are positively correlated in the near-diploid cell lines, but not in the highly-aneuploid cell lines. Pearson’s correlation (N=261 for ANAPC4 vs CDC20 mRNA expression in the bottom aneuploidy quartile, N=270 for ANAPC4 vs CDC20 mRNA expression in the top aneuploidy quartile, N=96 for ANAPC4 vs CDC20 protein expression in the bottom aneuploidy quartile, N=97 for ANAPC4 vs CDC20 protein expression in the top aneuploidy quartile) **(C,D)** Quantification of cell viability of the highly-aneuploid derivatives of HCT116,HPT1 **(C)** and HPT2 **(D)** with and without CDC20 (left) or ANAPC4 (right) knockout, following SAC inhibition (125 nM reversine). CDC20 depletion in aneuploid cells significantly reduced sensitivity to SAC inhibition, while ANAPC4 depletion did not. One way ANOVA (N = 3; *, p-value<0.05; **, p-value<0.01).

To validate the role of the increased CDC20 levels in the increased sensitivity of aneuploid cells to SAC inhibition, we knocked out CDC20 or ANAPC4 in two highly-aneuploid derivatives of HCT116, HPT1 and HPT2 (Kuznetsova *et al*, 2015, **Supplementary Fig. 3**), and assessed their cellular sensitivity to reversine. In both aneuploid cell lines, CDC20 knockout decreased the cellular sensitivity to SAC inhibition, whereas ANAPC4 knockout did not have the same effect (**Figure 3C,D**).

### CDC20 expression levels determine the prevalence of mitotic errors and metaphase duration

Next, we set out to mechanistically explore the association between aneuploidy, CDC20 expression levels, and the response to SAC inhibition. We quantified CDC20 protein levels in two isogenic cellular systems of near-diploid human cell lines – the colon cancer cell line HCT116, and the non-transformed retinal cell line RPE1 – and their highly-aneuploid derivatives (Kuznetsova et al, 2015). As CDC20 protein levels fluctuate throughout the cell cycle (Foe et al, 2011), they were quantified at their peak, during metaphase, in dividing single cells using immunofluorescence microscopy. In both systems, highly-aneuploid cells expressed significantly higher levels of CDC20 compared to their near-diploid counterparts (**Figure 4A** and **supplementary Fig. 4A**). Importantly, this over-expression of CDC20 during mitosis in individual cells indicates that the elevated expression of CDC20 that we previously observed in cell populations does not merely reflect changes in their proliferative capacity, but rather changes at the single cell level. We previously found highly-aneuploid cells to exhibit more mitotic errors and linked their CIN to their elevated response to SAC inhibition (Cohen-Sharir et al, 2021). We therefore hypothesized that the elevated CDC20 expression levels may be associated with the elevated CIN of the aneuploid cells.

**Figure 4.**
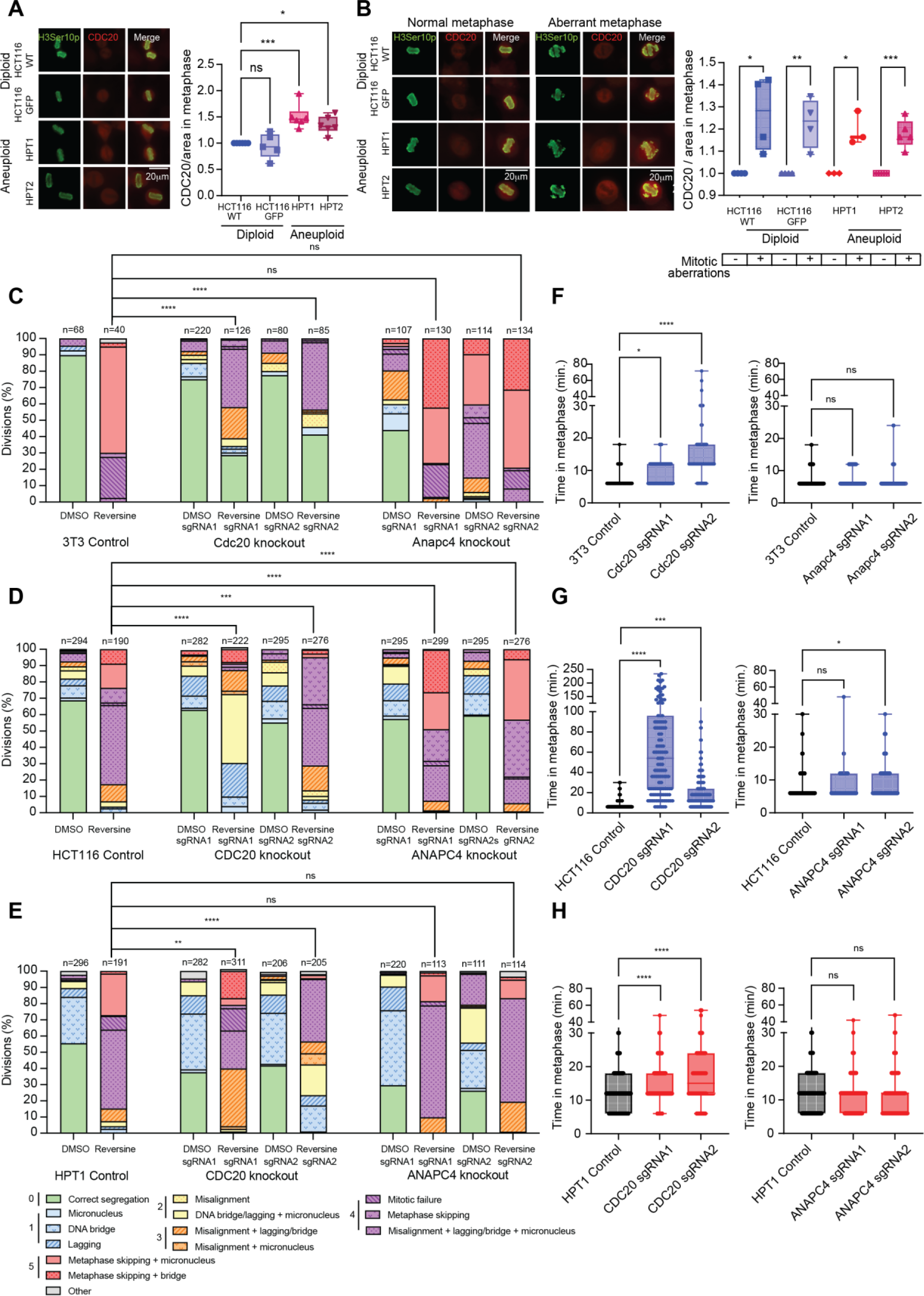
CDC20 expression levels determine the prevalence of mitotic errors and metaphase duration. **(A,B)** Representative immunofluorescence images (left) and quantification (right) of CDC20 protein levels during metaphase, in single HCT116 cells and their highly-aneuploid derivatives. Cells were synchronized to the G2/M border using RO-3306, released from arrest, fixed upon arrival to metaphase and stained for CDC20 and for the mitosis marker H3Ser10p. **(A)** Highly-aneuploid HPT cells express higher levels of CDC20 than their diploid counterparts. One-way ANOVA (N=6, *, p-value <0.05; ***, p-value <0.001). **(B)** Imaging (left) and quantification (right) of CDC20 in cells undergoing normal or aberrant mitoses. Cells with mitotic aberrations express significantly higher levels of CDC20 during metaphase than cells undergoing normal division, regardless of ploidy background. Two-sided t-test (N=5; *, p-value<0.05; **, p-value<0.01; ***, p-value<0.001). **(C-E)** Quantification of mitotic abnormalities in cells treated with 125nM (or 250nM) reversine, under control conditions (left), CDC20 knockout (middle) or ANAPC4 knockout (right). Mitotic aberrations were identified by live-cell imaging and scored on a severity scale of 0-5, then grouped and colored by score. The changes in severity distribution across samples were assessed using one-sided Kruskal Wallis tests, as elaborated in the Methods section. NIH-3T3 **(C)** and HPT1 **(E)** cells treated with 250nM or 125nM reversine respectively exhibit reduced mitotic aberrations after CDC20 knockout (middle), but not after ANAPC4 knockout (right). One-sided Kruskal Wallis (****, p-value < 0.0001). **(D)** HCT116 cells treated with reversine exhibit decreased mitotic aberrations after CDC20 knockout (middle) and increased mitotic aberrations after ANAPC4 knockout (right). One-sided Kruskal Wallis (*, p-value < 0.05; ****, p-value < 0.0001). **(F-H)** Time in metaphase for indicated genotypes quantified by live-cell imaging. CDC20 knockout (left) prolongs time in metaphase in NIH-3T3 **(F)**, HCT116 **(G)** and HPT1 **(H)** cells. ANAPC4 knockout (right) does not significantly affect time in metaphase in NIH-3T3 **(F)** and HPT1 **(H)** cells and shortens metaphase duration in HCT116 cells **(G)**. One-way ANOVA, (N= 315 for 3T3-Cdc20 knockout, N= 194 for 3T3-Anapc4 knockout, N= 496 for HCT116-CDC20 knockout, N= 666 for HCT116-ANAPC4 knockout, N=608 for HPT1-CDC20 knockout, N=586 for HPT1-ANAPC4 knockout; *, p-value<0.05; **, p-value<0.01; ***, p-value<0.001; ****, p-value < 0.0001).

To test this, we compared CDC20 protein expression in cells undergoing normal and erroneous cell divisions and found that in both isogenic systems – regardless of ploidy background – cells undergoing aberrant mitosis presented higher levels of CDC20 during metaphase (**Figure 4B** and **Supplementary Fig. 4B**). These results suggest a link between increased CDC20 expression and chromosome missegregation, which is both a cause and a consequence of the aneuploid state (Pfau & Amon, 2012; Ben-David & Amon; Holland & Cleveland, 2009).

To test whether CDC20 expression was directly linked to SACi-induced CIN, we characterized the effect of CDC20 levels on the overall rate and severity of mitotic errors in mouse and human cells using live cell imaging. To this end, we quantified mitotic aberrations in control and CDC20 knockout cells, with or without SAC inhibition, and scored the mitotic aberrations on a severity scale of 0-5, with 0 (green) representing normal division and increasingly higher scores (warmer colors) representing more severe mitotic aberrations, as has been done in previous works (Crozier et al, 2022; Thu et al, 2018; Huis In ’T Veld et al, 2019; see Methods section). Both in NIH-3T3 and HCT116 cells subjected to SAC inhibition, CDC20 knockout significantly reduced the prevalence and the severity of reversine-induced mitotic aberrations, thus alleviating SAC-inhibition-induced CIN (**Figure 4C** and **Figure 4D**, respectively). Similarly, CDC20 knockout alleviated CIN in the highly-aneuploid HPT1 and HPT2 cells (**Figure 4E** and **Supplementary Fig. 4C**, respectively). In contrast, ANAPC4 knockout in NIH-3T3 (**Figure 4C**), HPT1 (**Figure 4E**) and HPT2 (**Supplementary Fig. 4C**) did not significantly affect the prevalence of mitotic aberrations following SAC inhibition. In HCT116 cells, ANAPC4 knockout even led to an increase in the severity of mitotic aberrations (**Figure 4D**). Together, these findings demonstrate a causal relationship between CDC20 expression, and the degree of CIN induced by SAC inhibition. Specifically, reduction in CDC20 levels is associated with decreased severity and overall rate of mitotic aberrations in mouse and human cells of various transformation and ploidy statuses.

Low expression and activity of CDC20 are expected to prolong mitosis, which might underlie the association that we observed between CDC20 and CIN levels. We thus quantified metaphase duration following gene knockout in all cell lines. CDC20 knockout significantly prolonged metaphase duration in NIH-3T3 (**Figure 4F**), HCT116 (**Figure 4G**), HPT1 (**Figure 4H**) and HPT2 **(Supplementary Fig. 4D**) cells. In contrast, all cell lines except HCT116 did not present a significant change in metaphase duration after ANAPC4 knockout (**Figure 4F-H** and **Supplementary Fig. 4D**). The differences in mitotic phenotypes and duration in the NIH-3T3 and HCT116 cell lines with and without CDC20 and ANAPC4 knockout can be observed in **Supplementary Videos 1–10**. All in all, CDC20 depletion, but not depletion of the APC/C subunit ANAPC4, results in prolonged metaphase and reduced prevalence of mitotic aberrations across mouse and human cell lines with various ploidies. These results indicate that CDC20 is the key player in the cellular response to SAC inhibition, and that it acts by altering metaphase duration and affecting the overall level of chromosomal instability induced by SAC inactivation.

## Discussion

Aneuploidy and CIN are associated with cancer progression and drug response. CIN and aneuploidy were previously linked to increased drug resistance to many anticancer drugs (Lee et al, 2011; Replogle et al, 2020; Ippolito et al, 2021; Lukow et al, 2021; Cohen-Sharir et al, 2021), but they can also lead to increased sensitivity to specific therapies, such as SAC inhibition (Cohen-Sharir et al, 2021), KIF18A inhibition (Cohen-Sharir et al, 2021; Marquis et al, 2021), Src1 inhibition (Schukken et al, 2020), IL-6R inhibition (Hong et al, 2022), MAPK signaling inhibition (Zerbib et al, 2023) and proteasome inhibition (Ippolito et al, 2023). Understanding the molecular mechanisms that mediate the associations between CIN/aneuploidy and drug response can guide the development of new therapies and promote our basic understanding of cancer.

SAC inhibitors operate by inactivating the chromosome segregation-control mechanism, culminating in a cascade of chromosomal instability and aneuploidy that ultimately leads to cellular death. Biomarkers that enable predicting which patients respond well to SAC inhibitors are urgently needed. Previous studies have suggested that the response to SAC inhibition is driven by the APC/C (Sansregret et al, 2017, Thu et al, 2018, Wild et al, 2016). Here, we show that the response is mostly driven by the expression and function of the main APC/C activator during mitosis, CDC20, rather than by the APC/C itself.

We show that CDC20 expression, but not APC/C expression, is significantly correlated to sensitivity to multiple forms of SAC inhibition and to the aneuploid state. Furthermore, we show that CDC20 depletion decreases the sensitivity to SAC inhibition in both mouse and human cells, while depletion of a representative APC/C subunit does not. Mechanistically, we show that CDC20 overexpression is significantly associated with mitotic errors, while CDC20 depletion results in prolonged metaphases and decreased prevalence and severity of mitotic errors under SAC inhibition. These results suggest that cells with lower CDC20 levels delay anaphase onset and spend more time in metaphase, allowing for more time to correct spindle abnormalities. This would lead to the observed decrease in mitotic error rate and severity, to the acquisition of more fit karyotypes and to the overall better survival of the cells under SAC inhibition.

Our study provides a molecular link between aneuploidy and the sensitivity to SAC inhibition. We show for the first time that aneuploid cells overexpress CDC20 and that this upregulation is correlated with an increased rate of mitotic aberrations, presumably leading to increased inherent chromosomal instability in these cells and to their enhanced sensitivity to the excess CIN caused by SAC inhibition. Therefore, the upregulation of CDC20 in aneuploid cells could be important for their adaptation to the aneuploid state, and at the same time mediate their increased sensitivity to SAC inhibition.

Effectively, SAC inhibition and CDC20 overexpression result in a similar outcome – they both lead to higher effective concentrations of uninhibited APC/C-CDC20 complexes in the cells, allowing cell division to continue even in the presence of malformed spindles. Therefore, each of these two conditions individually leads to chromosomal instability and aneuploidization (Weaver & Cleveland, 2009; Mondal et al, 2007; Mason et al, 2017; Adell et al, 2023). Combined, we show that these conditions lead to excessive chromosomal instability and the formation of unfit karyotypes, which is poorly tolerated by the cells. In contrast, CDC20 downregulation reduces the overall levels of chromosomal instability in the cells, allowing them to better tolerate the effects of SAC inhibition, thereby making them more resistant to this class of drugs.

CDC20 overexpression is known to be a negative prognostic marker in multiple cancer types (Xian et al, 2022; Wu et al, 2013; Karra et al, 2014; Kato et al, 2012), possibly due to its association with higher cell division rate of tumors (Shang et al, 2018; Chu et al, 2019). Here, we show that CDC20 overexpression might be a favorable prognostic marker of the response to SAC inhibitors, and perhaps even more so in highly-aneuploid tumors. Our findings are highly clinically relevant as several SAC inhibitors are currently in clinical trials, with one of them being fast-tracked by the FDA (NCT05251714). In a broader context, CDC20 may be a potential biomarker for predicting the cellular response to other CIN-inducing therapies, which needs to be validated in follow-up work.

## Materials and Methods

### CRISPR-Cas9 screen and Validation

NIH-3T3 cells were transduced with a doxycycline-inducible Cas9 system using lentiviruses. For this, 293FT cells were co-transfected with the Lenti-iCas9-neo vector (Addgene plasmid #85400, a gift from Qin Yan) (Cao et al, 2016), along with PMD2 and VSVG lentiviral vectors. Following transfection, cells were selected with 400μg/ml neomycin and monoclonal Cas9-expressing cells were isolated in 96-well plates. Cas9 activity was confirmed through the application of sgRNAs (IDT product #1072542) as positive controls, along with the T7 Endonuclease I (T7EI) assay (NEB).

To introduce the sgRNA library, 377 million iCas9-expressing NIH-3T3 cells were transduced with the Mouse Two Plasmid Activity-Optimized CRISPR Knockout Library (Wang et al, 2017) (Addgene; 1000000096) using spinfection at an MOI of 0.3 to ensure a final library coverage of 500X. The required number of iCas9 NIH-3T3 cells (377 million) was calculated as follows: the required minimal complexity (500X) multiplied by library complexity (188,509 sgRNAs) divided by minimal surviving fraction of cells at an MOI of 0.3 (25%).

Cells were then selected with puromycin for 5 days and frozen or maintained at a minimal complexity of 500X (94 million cells after puromycin selection). To identify sgRNAs enriched under reversine treatment, cells were expanded and seeded at 94 million cells per replicate (2 replicates per condition) with either DMSO (0.1%; Sigma) or 250 nM reversine (R3904; Sigma-Aldrich) on 15 cm dishes (833903; Sarstedt) and cultured for 5 days with treatment after which the treatments were removed.

Cells were then maintained and passaged for three more weeks when necessary (DMSO-treated cells ∼every 3 days, and reversine-treated cells once or twice at later time points), always ensuring a minimal number of 94 million cells per replicate. The whole screen (2 conditions, 2 replicates per condition) was performed twice with a small variation between screens: in the first iteration, cells were treated with doxycycline to induce iCas9 directly following transduction, while in the second iteration doxycycline was added 5 days after DMSO or reversine were added to the cultures. At the end of the screen, DNA was isolated from a minimum of 94 million cells for each replicate to determine sgRNA distribution.

Genomic DNA was isolated from samples using the QIAamp DNA Blood Maxi Kit (Qiagen; 56404). DNA was fragmented using Ndel and SacII restriction enzymes (New England Biolabs) and hybridized overnight with biotinylated capture oligos for selective DNA targeting. The hybridized DNA was captured using Streptavidin T1 Dynabeads (ThermoFisher; 65601) and non-hybridized oligos were removed by Exonuclease I digestion. sgRNAs were amplified by two rounds of PCR. The first round used unique barcoded forward primers, allowing for sample identification in pooled sequencing, while the second round attached necessary adapter sequences for sequencing compatibility. PCR products were purified using the NucleoSpin Gel and PCR Clean-up kit (Machery-Nagel; 740609.50). Sequencing was performed on a NextSeq 500 sequencer (Illumina).

sgRNA enrichment was determined using DESeq2, followed by the MAGeCK ranking aggregation method (Robust Rank Algorithm (RRA), (Love et al, 2014; Li et al, 2014) with multiple testing corrections by the Benjamini–Hochberg method. Our protocol was developed by adapting and refining methodologies from the study by Vera E. van der Noord *et* al (van der Noord et al, 2023) and can be found in the Supplementary information.

#### Computational Analysis of Sequencing Data

The raw sequencing data (fastq files) were first sorted into separate files using sample-specific barcodes (the first six bases of the read sequences contained the barcodes; Perl script). Another Perl script was subsequently used to obtain the sgRNA counts from the individual (sample-specific) fastq files. This script searches for a key sequence (CGAAACACC), which precedes the sgRNA sequences. This key sequence was detected within a certain window (bases 16-44). When found, it extracts the sgRNA sequences which are the 20 bases just after the key sequence or the 20 bases after a G base that sometimes follows the key sequence. When one of the two sequences matched with the list of known sgRNA sequences, the count of this sgRNA was raised by one. The sgRNA sequences were obtained from the Addgene website (Wang et al, 2017), https://www.addgene.org/pooled-library/sabatini-crispr-mouse-high-activity-two-plasmid-system/). Next, a differential analysis was performed on the sgRNA level (counts) between the two conditions (reversine and DMSO) with DESeq2 (Love et al, 2014). A paired design was implemented to account for differences between the samples while estimating the effect due to the condition (design=∼ replicate + condition). The median of ratios method was used for the normalization of the sgRNA counts. This analysis resulted in a log2 fold change value, a p-value, and a test statistic for each sgRNA. sgRNAs with a zero count in all samples were excluded. The results were subsequently ranked on the test statistic putting either the most enriched or depleted sgRNA on top. MAGeCK’s RRA tool (Kolde et al, 2012) was subsequently used to find genes that were consistently ranked better than expected under the null hypothesis of uncorrelated inputs (Settings: --permutation 100; -p [maximum percentile] = proportion of sgRNAs with positive or negative test statistic [depending on what is tested; positive or negative selection] and p-value <= 0.25). MAGeCK’s RRA tool uses the Benjamini-Hochberg method for multiple testing corrections (Benjamini & Hochberg, 1995).

### Computational analyses and statistics

For analysis of gene set enrichment within the CRISPR screen ranking results, enrichment was measured using the original GSEA method (Subramanian et al, 2005) (based on the estimated log-fold-change), which estimates the concentration of each gene set in the list of up-and down-regulated genes. We used the GSEA implementation in the GSEA-MSigDB website (https://www.gsea-msigdb.org/gsea/msigdb) to identify pathways enriched in the top 10% ranking genes of both CRISPR screens. The collections of gene sets used were the “KEGG”, “Hallmark” and “GOBP” gene set collections from MSigDB v.2023.2.Mm (Liberzon et al, 2011).

For analysis of the top genes enriched in both CRISPR screens, the top 25% of genes of each screen were intersected and their significance was compared.

mRNA and protein expression datasets were obtained from DepMap release 22Q2 (Tsherniak et al, 2017). SAC genetic dependency data were obtained from the Achilles genome-wide RNAi screen (release 22Q2), and drug sensitivity data were obtained from the PRISM repurposing primary screen (release 19Q4), both available on the cancer DepMap. For single-gene mRNA expression analyses, mRNA expression Z-score were calculated. For multiple-gene mRNA expression analyses, a gene set enrichment analysis (ssGSEA) score was assigned for each signature. Aneuploidy scores (AS) for all cell lines were obtained from Cohen-Sharir et al, Nature (2020) (Cohen-Sharir et al, 2021). General analyses were performed on the ∼1,700 cell lines documented in the DepMap. Ploidy-centered analyses were performed on a subset of ∼1,000 cells for which an aneuploidy score was previously calculated.

The DepMap cancer cell lines were split into two groups of near-diploid and highly-aneuploid cell lines, correlating to the quartiles with the bottom and top aneuploidy scores. Two-sided t-tests were used to compare gene expression between the groups. The cells were also split into groups of the top and bottom CDC20 or subunit-only-APC/C expression quartiles, and two-sided t-tests were used to compare drug sensitivity between the groups. Statistical analysis for viability/proliferation, and comparison of fluorescence intensity levels, between multiple groups were performed either by two-sided t-tests (when two groups were compared) or by one-way ANOVA (when more than two groups were compared). All statistical analyses were performed in GraphPad Prism 9.1.

### Tissue culture

NIH-3T3 and 293FT cell lines were procured from the American Type Culture Collection (ATCC). RPE1 and HCT116 cells, and their aneuploid derivatives HPT1, HPT2, RPT1, RPT3, and RPT4 included in this study were generated in Kuznetsova et al (Kuznetsova et al, 2015). All cells were cultured in Dulbecco’s Modified Eagle Medium (DMEM; Gibco; 31966-021) supplemented with 10% fetal bovine serum (FBS; Thermo Fisher Scientific or Sigma-Aldrich; 11573397), 100 U/mL penicillin and 100 U/mL Streptomycin (P/S; Gibco; 15140-122). Cells were grown at 37°C and 5% CO2. For passaging and subculturing, cells were detached using either 0.25% Trypsin-EDTA (Life Technologies) or Tryple Express (Gibco; 12605-010).

### Lentiviral transduction

To produce lentiviruses, 293FT cells were transfected with 3µg of the selected vector, complemented with essential packaging plasmids: 3µg of pSPAX2 and 1µg of pMD2.G. Notably, pSPAX2 (Addgene plasmid #12260) and pMD2.G (Addgene plasmid #12259), both gifts from Didier Trono. 48 hours after transfection, the medium from the 293FT cells was harvested, filtered through a 0.45 µm filter (VWR Science), and then directly added to the intended cells in the presence of 8-10 µg/ml polybrene (TR-1003-G, Sigma-Aldrich).

### CRISPR-Cas9 mediated gene knockout and shRNA mediated knockdown

For CRISPR knockout cell lines, sgRNAs targeting mouse and human genes of interest (see **Supplementary Table 2**) were cloned into plenti-sgRNA (Addgene plasmid #71409) using BsmbI and into Cas9 plasmid (Addgene plasmid #62988)(Ran et al, 2013) using BbsI enzyme (NEB). The vectors, including Lenti-iCas9-neo (Addgene plasmid #85400) (Cao et al, 2016) was a gift from Qin Yan, and the pSpCas9(BB)-2A-Puro V2.0 (PX459) was a gift from Feng Zhang (Addgene plasmid #62988) (Ran et al, 2013). For NIH-3T3 cell line knockouts targeting Cdc20 and Anapc4 genes, a combination of Lenti-iCas9-neo and plenti-guideRNA vectors were used (see lentiviral transduction section below), followed by selection with 400 µg/ml G418 (Thermofisher) for 2 weeks, 1 µg/ml Puromycin (Invitrogen) for 5 days, and 1 µg/mL Doxycycline induction. In HCT116, HPT1, and HPT2 cell lines, CDC20 and ANAPC4 knockouts were obtained with sgRNA plasmids transfected using FuGENE® HD Transfection Reagent (Promega, E2311), followed by selection using puromycin (Invitrogen, 0.1-1µg/ml) for 5 days.

shRNAs were cloned into the Tet-pLKO-puro vector (Addgene plasmid #21915) (Wiederschain et al, 2009) using AgeI and EcoRI restriction enzymes (NEB) and verified by sequencing. Following lentiviral transduction, target cells were selected with puromycin (3-7 days) to isolate successfully transduced cells. Inducible shRNA expression was activated using 1 µg/mL doxycycline (hyclate D9891, Sigma Aldrich) in the culture medium. Gene knockdown efficacy was confirmed by quantitative RT-PCR post three-day induction, and gene knockout effectiveness was assessed via western blotting. The cellular impact of gene silencing was evaluated using various assays, including a five-day crystal violet staining protocol.

### Time-lapse imaging

Chromosomal abnormalities were quantified using time lapse imaging. For this, cell lines were transduced with lentiviral H2B-mCherry constructs. One day prior to imaging, 3.2×105 NIH-3T3 cells, 6.8×105 HCT116 cells, and 3.2×105 cells for both HPT1 and HPT2 were seeded into imaging disks (Greiner Bio-One, #627870). Imaging was performed on a DeltaVision Elite microscope (GE Healthcare), fitted with a CoolSNAP HQ2 camera and a 40X, 0.6 NA immersion objective lens (Olympus) for at least 20 hours with images captured every 6 minutes. Each imaging stack consisted of 30 to 40 Z-stacks, at 0.5 µm apart. Images were analysed using ICY software (Institut Pasteur). Only mitotic cells were included in the analyses. For live-cell imaging, the cells were pretreated with the drug one hour before initiating the imaging sessions.

### Scoring System for Chromosomal Instability Resulting from Missegregation

To quantify CIN phenotypes, mitotic phenotypes were scored according to previously defined standards (Crozier et al, 2022; Musacchio & Salmon, 2007; Thu et al, 2018). A minority of chromosomal events, representing diverse chromosomal aberrations and anomalies that did not align with standard categories were classified as ‘other’. Each of these ‘other’ events were scored from 1 to 5, based on its severity and complexity, relative to the predefined categories. However, due to their atypical nature, ’other’ events were excluded from the main statistical analysis. The missegregation events in **Figure 4C-E** and **Supplementary Fig. 4** are colored by score, with specific events within the same scoring category marked with different patterns.

<u>points (green):</u> Correct chromosomal Segregation.

<u>point (blue):</u> DNA Bridge Formation, Micronucleus Formation, Chromosomal Lagging.

<u>points (yellow):</u> DNA Bridge/Lagging with Micronucleus Formation, Metaphase Misalignment.

<u>points (orange):</u> Metaphase Misalignment with Micronucleus Formation, Metaphase Misalignment with Chromosomal Lagging/Bridge Formation.

<u>points (purple):</u> Metaphase Skipping, Mitotic Failure, Metaphase misalignment with chromosomal lagging/bridge formation and emergence of micronuclei.

<u>points (red):</u> Metaphase Skipping with DNA Bridging, Metaphase Skipping with Micronucleus Formation.

<u>Other (Variable 1-5 points, grey):</u> Chromosomal aberrations or anomalies that do not fit into the predefined categories.

For each sample, aberrations of categories 0-5 were grouped by score, and the differences in their distribution across samples were calculated using one-sided Kruskall-Wallis tests performed in GraphPad Prism 9.1.

### Assessment of cell viability using crystal violet staining

For crystal violet viability assays, cells were seeded in 12-well plates at 10-20% confluency and treated with DMSO or 125nM (or 250nM) reversine for five days. Cells were fixed in 4% paraformaldehyde for 15-20 minutes at room temperature. Cells were washed with PBS and stained with 0.1% w/v crystal violet for 8-10 minutes. Excess dye was washed off with distilled water, and plates were air-dried. For quantification, crystal violet was extracted with 0.5-2 ml of 10% acetic acid. Absorbance was measured in 96 well plates at 570-595 nm using a microplate reader (Thermo multiskan Go).

### Quantification of proliferation using EdU incorporation

For EdU assays, cells were cultured on coverslips in 24-well plates and treated with 125 nM (or 250nM) Reversine or DMSO for 48 hours. To determine the fraction of cells in S-phase, cells were pulse-labelled with 10 µM EdU for 2 hours. Cells were fixed in 4% formaldehyde and EdU was detected using Click chemistry and (2 mM Cu(II)SO₄, 4 µM sulfo-Cy3-azide, and 20 mg/ml sodium ascorbate in PBS). EdU incorporation was visualized by fluorescence microscopy (Olympus IX51 or Olympus BX43), and images processed and quantified using the Fiji software (ImageJ 1.53C).

### Western blotting

For Western blotting, cells were lysed in ELB (150 nM NaCl, 0.1% NP-40, 5 nM EDTA, and 50 mM HEPES (pH 7.5)), supplemented with a Roche protease inhibitor cocktail for 30 minutes. 20-30 µg of protein was loaded on 7.5-10% polyacrylamide gels and transferred to PVDF membranes. Membranes were blocked with Odyssey blocking buffer or 5% milk in TBST. Primary antibodies used were CDC20 (Bethyl, A301-180A, 1:1000), β-Actin (Cell Signaling, 4970 or 3700S, 1:2000), and ANAPC4 (Cusabio, CSB-PA135409, 1:1000) with incubation overnight at 4°C. Secondary antibodies (IRDye 800CW Goat anti-Rabbit IgG (H+L) (Licor; 1:15,000) and IRDye 680CW Goat anti-Mouse IgG (H+L) (Licor; 1:15,000)) were incubated for 1 hour at room temperature. Blots were visualised on an Odyssey imaging system (LI-COR Biosciences), in combination with Image Studio Lite software (LI-COR Biosciences).

### Real time quantitative PCR

For qPCR analysis, RNA was extracted from cell pellets using the RNA plus isolation kit (Qiagen, MN 740984.250). Detailed sequences of the qPCR primers are provided in Supplementary Table 2. For cDNA synthesis, 1.5 μg of RNA was used with the LunaScript RT SuperMix Kit (Bioke, M3010X) in a 20 μl reaction. The synthesized cDNA was then quantified by quantitative PCR (qPCR) using iTaq Universal SYBR Green supermix (Biorad, 1725124) on a LightCycler® 480 Instrument. Relative RNA levels were calculated in Excel (Microsoft) and plotted using Prism software (GraphPad).

### Immunofluorescence microscopy

For immunofluorescence imaging, cells were grown on glass coverslips in 24 well plates at a density of 2×10^5^ (HCT-HPT) or 1.5×10^5^ (RPE-RPT) cells. Cells were synchronized at the G2/M transition with 9nM (HCT-HPT) or 4.5nM (RPE-PRT) of the CDK1 inhibitor RO-3306 (Sigma-Aldrich, SML0569) for 20-21 hours, released from arrest and incubated for 45-55 minutes at 37 °C. Cells were visually followed until metaphase by H2B-GFP staining in an Axio Imager Z microscope (Carl Zeiss) and fixed in 4% paraformaldehyde for 15 minutes. Fixed cells were incubated for 15 minutes with fresh 0.1M Glycine to prevent quenching, then for 5 minutes with a 0.5% Triton X-100 solution to permeabilize cells. Cells were blocked for 30 minutes (BSA, Glycine, NaCl and 0.1% Triton X-100) and stained with primary antibodies against CDC20 (1:50, Santa Cruz) and the mitosis marker H3ser10p (1:400, Cell Signaling) in blocking buffer. Used secondary antibodies were anti-mouse Alexa-555 antibody (1:400, Cell Signaling) and an anti-rabbit Alexa-488 (1:400, Cell Signaling). Both primary and secondary antibodies were incubated for 1 hour in a humidified environment at room temperature. Images were acquired using CellSens Imaging Software (Olympus) at a 40x resolution. Cells at metaphase were identified by H3ser10p staining and CDC20 intensity per area was quantified using ImageJ v1.53 in single cells. CDC20 levels were later compared in bulk between the isogenic cell lines in One-way ANOVA tests performed in GraphPad Prism 9.1.

## Acknowledgements

We are grateful to Roderick Beijersbergen and Cor Lieftink (Netherlands Cancer Institute, Amsterdam) for kindly providing scripts and advice for the analysis of the CRISPR screen data; and to Zuzana Storchova (RPTU Kaiserlautern, Germany) for kindly providing the HPT1, HPT2, RPT1, RPT3 and RPT4 cell lines. Siqi Zheng and Lin Zhou were supported by personal fellowships from the Chinese Scholar Council (CSC). This work was further supported by two Dutch Cancer Society grants to Foijer (2015-RUG-7822 and 2022-4EXPL-14805; ScreeninC). Work in the Ben-David Lab was supported by the DoD CDMRP Career Development Award (grant #CA191148 to U.B.-D.), the European Research Council Starting Grant (grant #945674 to U.B.-D.), the Israel Cancer Research Fund Project Award (U.B.-D.), the Azrieli Foundation Faculty Fellowship (U.B.-D.), the Israel Science Foundation (grant #1805/21 to U.B.-D.), the BSF Project Grant (grant #2019228 to U.B.-D.), and the Israel Cancer Association (grant #20230018 to U.B.-D.).

## Author contributions

SZ, LR, and LZ were involved in project planning and experimental design. FF and UB-D designed and supervised the study and provided funding. SZ, LR, LZ, YC-S and RT contributed to data acquisition, processing, and analysis. RT, MS-PR, SW, AvdB, performed live-cell imaging experiments. MB prepared CRISPR screen libraries, RW performed the CRISPR screen computational processing. PB was involved in data acquisition. SZ and LR wrote the manuscript together with FF and UB-D. All authors proofread the manuscript.

## Dataset availability

All datasets generated and analyzed in the CRISPR screen are available in the European Nucleotide Archive (ENA) under accession ID PRJEB71335. This includes detailed sample data for our CRISPR screen in NIH-3T3 cells, using a sgRNA library for sgRNA enrichment analysis. The availability of these datasets ensures transparency and facilitates replication of our findings, adhering to the open science principles of the EMBO Journal.

## Supplementary Information

### Supplementary Tables

**Supplementary Table 1:** Complete gene rankings in both screens.

### Supplementary Movies

Videos depicting mitotic processes in NIH 3T3 wild-type **(1)**, NIH 3T3 Cdc20 sgRNA1 knockout (ko) **(2)**, and NIH 3T3 Cdc20 sgRNA2 knockout **(3)** cells, illustrating the cellular response to Cdc20 depletion. Visual representation of mitotic activity in **(4)** NIH 3T3 ANAPC4 sgRNA1 knockout **(4)** and NIH 3T3 Anapc4 sgRNA2 knockout **(5)** cells providing insight into mitotic alterations following Anapc4 gene knockout.

Videos depicting mitotic processes in HCT116 wild-type (WT) **(6)**, HCT116 CDC20 sgRNA1 knockout (ko) **(7)**, and HCT116 CDC20 sgRNA2 knockout **(8)** cells, illustrating the cellular response to CDC20 depletion. Visual representation of mitotic activity in HCT116 ANAPC4 sgRNA1 knockout **(9)** and HCT116 ANAPC4 sgRNA2 knockout **(10)** cells, providing insight into mitotic alterations following ANAPC4 gene knockout.

### Legends to supplementary figures

**Supplementary Figure 1.**
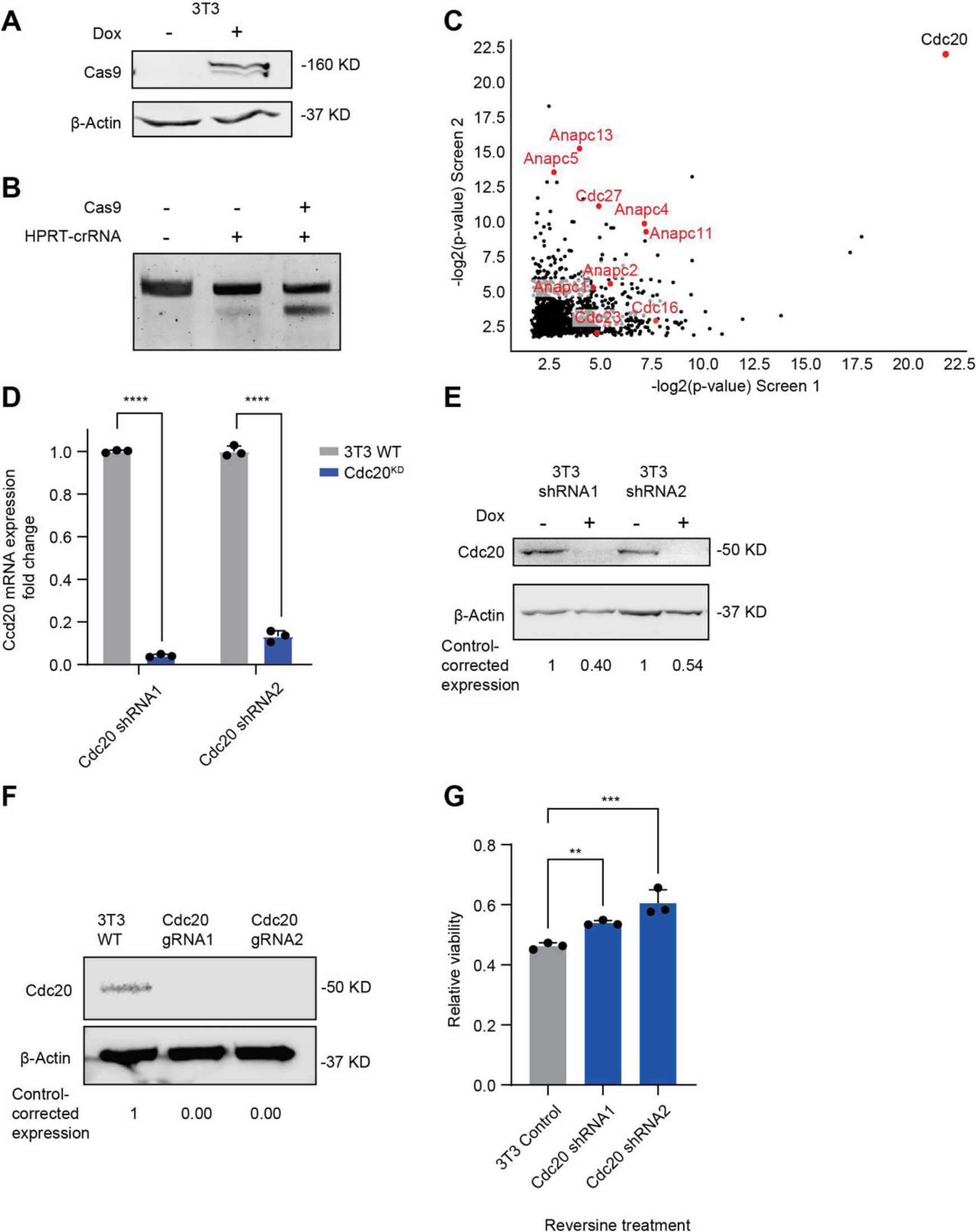
Cdc20 is strongly associated with the response to SAC inhibition. **(A)** Western blot for Cas9 expression in 3T3 cells used for the CRISPR screen. **(B)** Cleaved and uncleaved PCR product in a T7 assay as a readout of Cas9 activity in 3T3 cells used for the CRISPR screen. **(C)** Correlation between the top ranked 25% of genes in both CRISPR screens, based on their statistical significance with all APC/C-related genes highlighted, showing that Cdc20 is by far the most significant outlier of all APC/C extended complex members. **(D)** qPCR validation of Cdc20 knockdown in 3T3 cells. Two-sided T-test (N=3; ****, p-value< 0.0001). **(E)** Cdc20 knockdown validation by qPCR. **(F)** Cdc20 knockout validation by Western blot. **(G)** Cdc20 depletion significantly reduced the sensitivity to SAC inhibition. One-way ANOVA (N = 3, **, p-value <0.01, ***, p-value <0.001; ****, p-value < 0.0001).

**Supplementary Figure 2:**
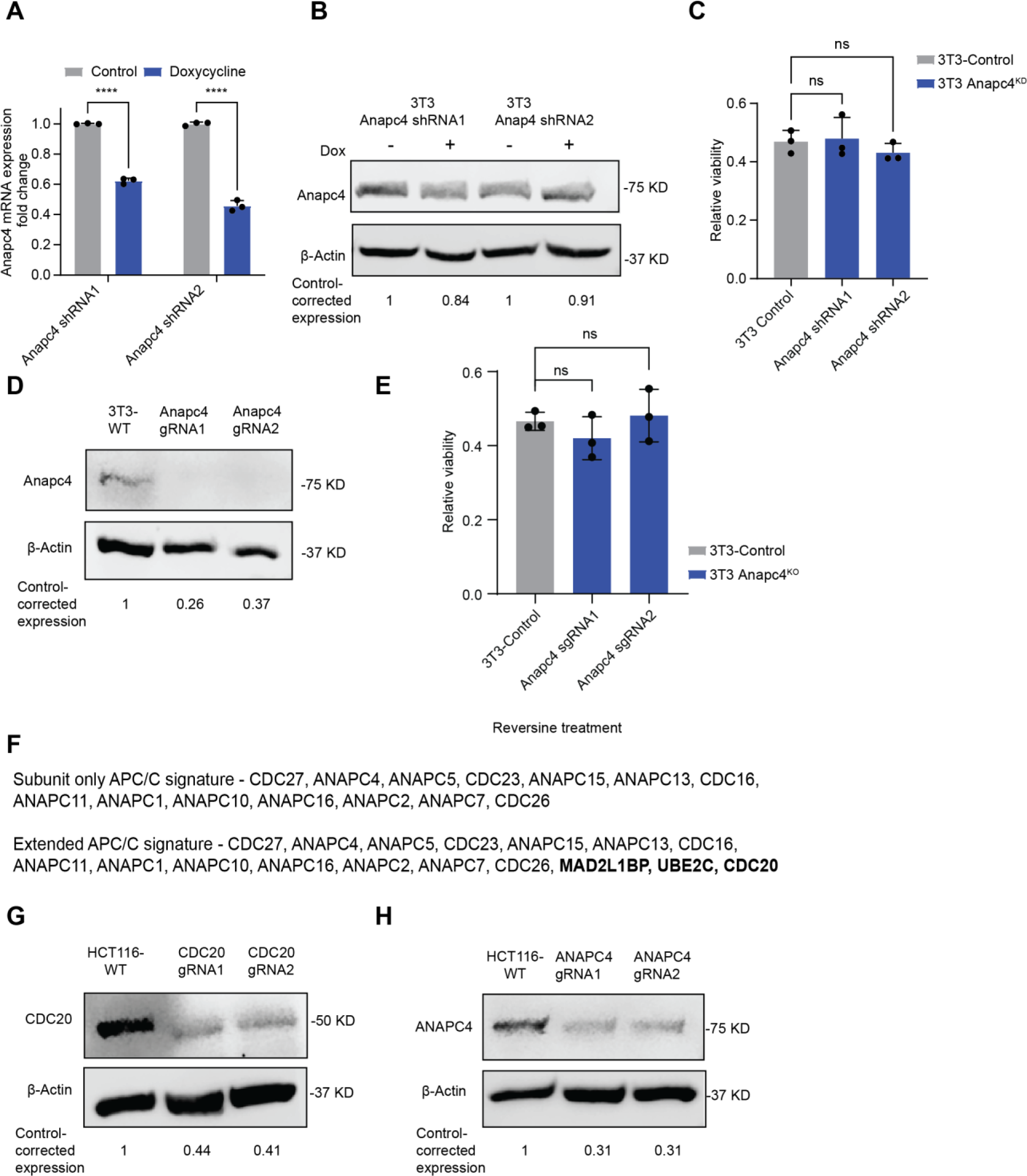
CDC20 expression, but not APC/C expression, predicts sensitivity to genetic and chemical SAC perturbation. **(A)** Anapc4 knockdown validation in 3T3 cells by qPCR. Two-sided T-test (N = 3; ****, p-value<0.0001). **(B)** Anapc4 knockdown validation in 3T3 cells by Western blot. **(C)** Viability of 3T3 cells treated with reversine with and without Anapc4 knockdown, relative to DMSO treated control cells. One-way ANOVA (N=3). **(D)** Validation of Anapc4 knockout in 3T3 cell by Western blot. **(E)** Viability of 3T3 cells treated with 250nM reversine, with and without Anapc4 knockout. Anapc4 knockdown does not change cell viability under Reversine. One-way ANOVA (N=3). **(F)** Gene signatures used to define the core and extended APC/C complex members. The “APC/C subunit-only” signature contains the 14 core APC/C subunits (Yamano, 2019), while the “extended APC/C signature” described by Thu et al (Thu et al, 2018) contains three additional APC/C co-factors, including CDC20. **(G-H)** Validation of CDC20 **(F)** and ANAPC4 **(G)** knockout in HCT116 cells by Western blot.

**Supplementary Figure 3.**
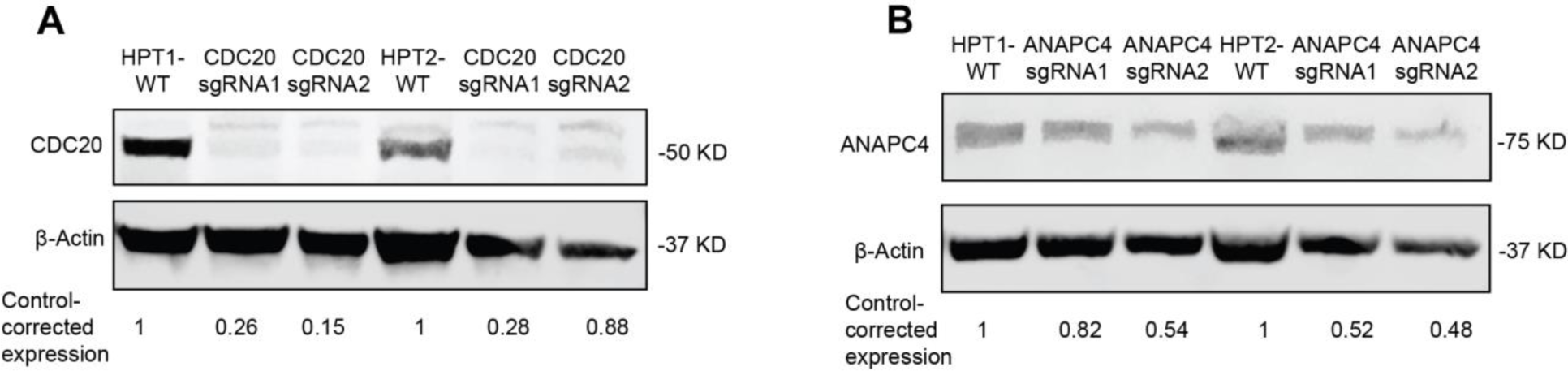
Knockout validation in aneuploid HPT cells. Shown are Western blot knockout validations for CDC20 **(A)** and ANAPC4 **(B)** in the HCT116 aneuploid derivatives HPT1 and HPT2.

**Supplementary Figure 4.**
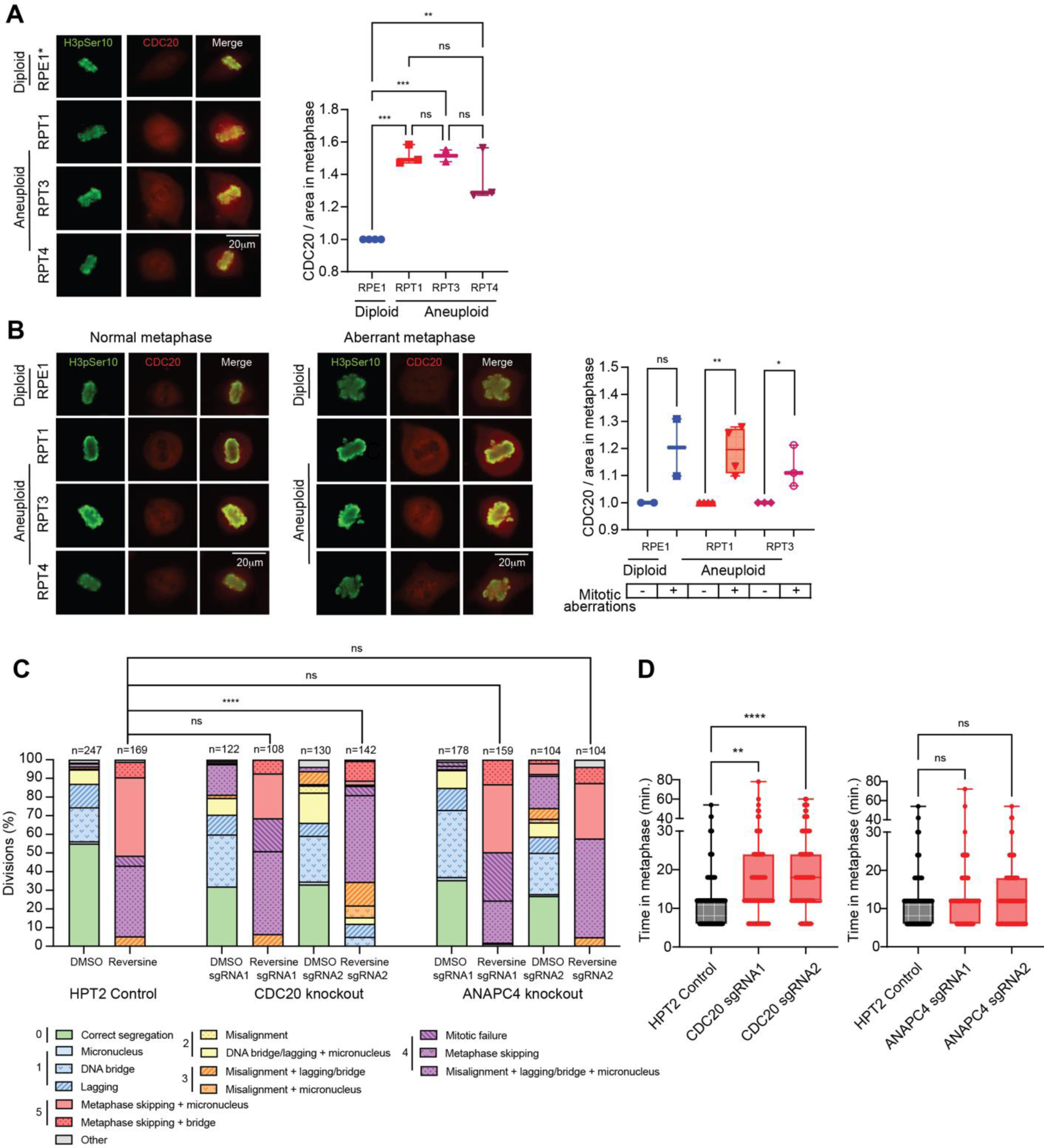
CDC20 expression levels determine the prevalence of mitotic errors and metaphase duration. **(A,B)** Representative IF images (left) and quantification (right) of CDC20 protein levels during metaphase, in single cells of RPE1 cells and their highly-aneuploid derivatives. Cells were synchronized to the G2/M border using RO-3306, released from arrest, fixed upon arrival to metaphase and stained for CDC20 and for the mitosis marker H3Ser10p. **(A)** highly-aneuploid cells express higher levels of CDC20 than their diploid counterparts. One-way ANOVA (N=4, **, p-value <0.01; ***, p-value <0.001). **(B)** Imaging (left) and quantification (right) of CDC20 in cells undergoing normal or aberrant mitoses. Cells with mitotic aberrations express significantly higher levels of CDC20 during metaphase than cells undergoing normal division, regardless of ploidy background. Two-sided t-test (N=5, *, p-value<0.05; **, p-value<0.01). **(C)** Quantification of mitotic abnormalities in HPT2 cells treated with 125nM reversine, under control conditions (left), CDC20 knockout (middle) or ANAPC4 knockout (right). Mitotic aberrations were identified by live-cell imaging, scored on a severity scale of 0-5, and then grouped and colored by score. The changes in severity distribution across samples were assessed using a one-sided Kruskal Wallis test, as elaborated in the Methods section. CDC20 knockout (middle) decreases the rate and severity of mitotic aberrations, while ANAP4 knockout (right) does not. One-sided Kruskal Wallis (****, p-value < 0.0001). **(D)** Time in metaphase of HPT2 cells with and without gene knockout, determined by live-cell imaging. CDC20 knockout (left) prolongs time in metaphase, while ANAPC4 knockout does not. One-way ANOVA (N=360 for HPT2-CDC20 knockout, N=406 for HPT2-ANAPC4 knockout; ****, p-value<0.0001).

**Supplementary Table 2:**
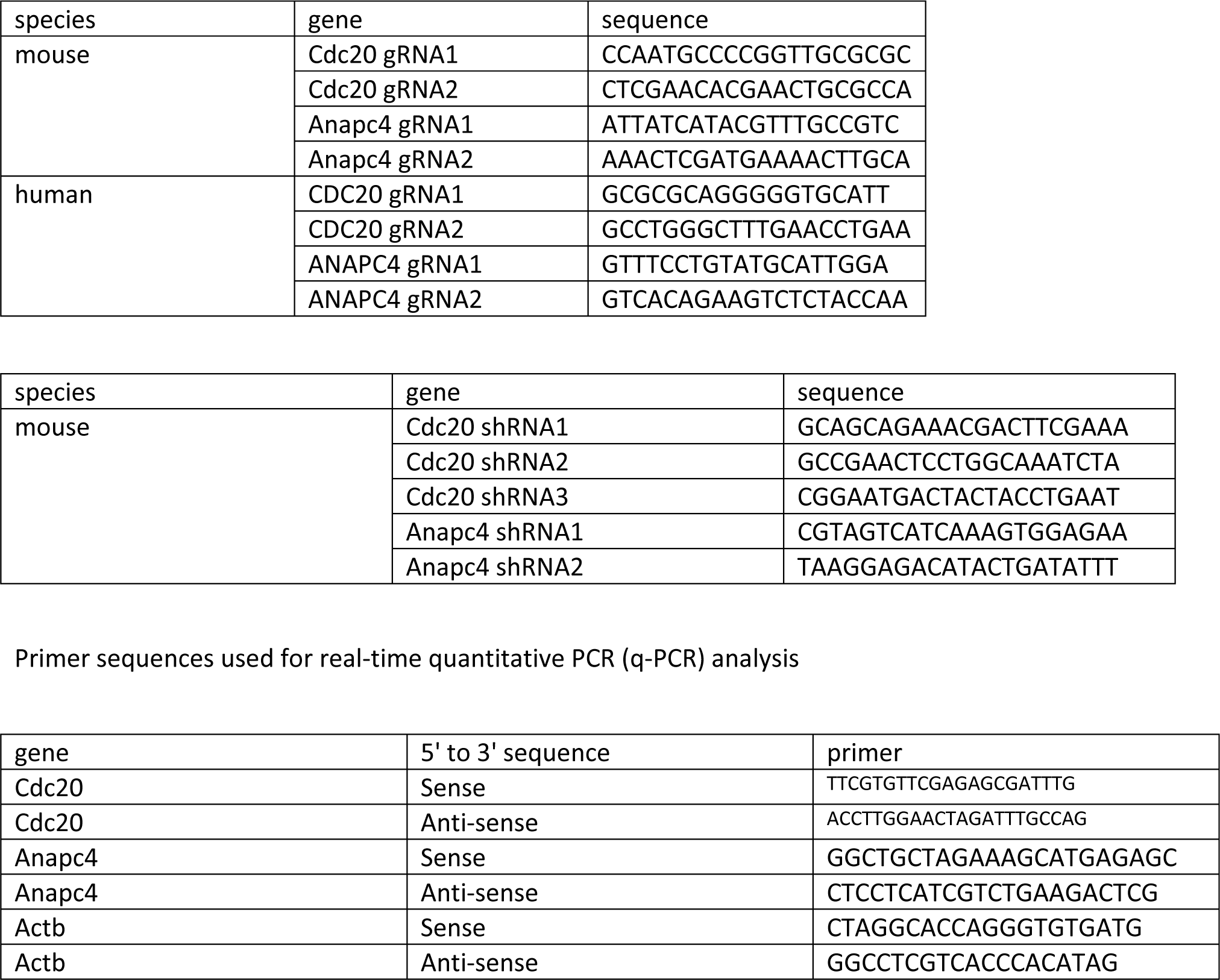
sgRNA and shRNA sequence data.

